# MORPHOLOGICAL DETERMINANTS OF CELL-TO-CELL VARIATIONS IN ACTION POTENTIAL DYNAMICS IN SUBSTANTIA NIGRA DOPAMINERGIC NEURONS

**DOI:** 10.1101/2021.12.07.471584

**Authors:** Estelle Moubarak, Yanis Inglebert, Fabien Tell, Jean-Marc Goaillard

## Abstract

Action potential (AP) shape is a critical electrophysiological parameter, in particular because it strongly modulates neurotransmitter release. AP shape is also used to distinguish neuronal populations, as it greatly varies between neuronal types. For instance, AP duration ranges from hundreds of microseconds in cerebellar granule cells to 2-3 milliseconds in substantia nigra pars compacta (SNc) dopaminergic (DA) neurons. While most of this variation seems to arise from differences in the subtypes of voltage- and calcium-gated ion channels expressed, a few studies suggested that dendritic morphology may also affect AP shape. However, AP duration also displays significant variability in a same neuronal type, while the determinants of these variations are poorly known. Using electrophysiological recordings, morphological reconstructions and realistic Hodgkin-Huxley modeling, we investigated the relationships between dendritic morphology and AP shape in SNc DA neurons. In this neuronal type where the axon arises from an axon-bearing dendrite (ABD), the duration of the somatic AP could be predicted from a linear combination of the complexities of the ABD and the non-ABDs. Dendrotomy simulation and experiments showed that these correlations arise from the causal influence of dendritic topology on AP duration, due in particular to a high density of sodium channels in the somato-dendritic compartment. In addition, dendritic morphology also modulated AP back-propagation efficiency in response to barrages of EPSCs in the ABD. In line with previous findings, these results demonstrate that dendritic morphology plays a major role in defining the electrophysiological properties of SNc DA neurons and their cell-to-cell variations.

**SIGNIFICANCE STATEMENT:** Action potential (AP) shape is a critical electrophysiological parameter, in particular because it strongly modulates neurotransmitter release. AP shape (e.g. duration) greatly varies between neuronal types but also within a same neuronal type. While differences in ion channel expression seem to explain most of AP shape variation across cell types, the determinants of cell-to-cell variations in a same neuronal type are mostly unknown. We used electrophysiological recordings, neuronal reconstruction and modeling to show that, due to the presence of sodium channels in the somato-dendritic compartment, a large part of cell-to-cell variations in somatic AP duration in substantia nigra pars compacta dopaminergic neurons is explained by variations in dendritic topology.

## INTRODUCTION

The diversity of neuronal types in the mammalian brain is associated with a significant variation in action potential (AP) shape, such that AP properties are sometimes used to distinguish between neighboring neuronal populations (Bean, 2007; Seutin and Engel, 2010; Bucher and Goaillard, 2011; Ding et al., 2011; Goaillard et al., 2020). In particular, AP duration (often defined by the width at half-amplitude or half-width) varies from one hundred microseconds in rat cerebellum granule cells to 2-3 milliseconds in substantia nigra pars compacta (SNc) dopaminergic (DA) neurons (Bean, 2007; Carter and Bean, 2009; Delvendahl and Hallermann, 2016; Moubarak et al., 2019; Goaillard et al., 2020). These variations in AP duration are often associated with differences in firing patterns: for instance, fast-spiking neurons tend to have short action potentials (sub-millisecond) while cells with broad action potentials such as DA or cholinergic neurons tend to fire at lower frequencies (Lien and Jonas, 2003; Bean, 2007; Alle et al., 2009; Sengupta et al., 2010; Delvendahl and Hallermann, 2016). In addition, AP duration may vary in a same neuron, for instance when the neuron is firing repeatedly at high frequency (Ma and Koester, 1996; Geiger and Jonas, 2000; Lien and Jonas, 2003), modifying in turn neurotransmitter release and the amplitude of synaptic responses in the post-synaptic neuron (Geiger and Jonas, 2000). From a mechanistic point of view, the density of ion channels underlying the upstroke and decay of the AP (in particular sodium and potassium channels), and the gating properties of the specific subunits expressed are partly responsible for the observed variation in AP duration (Bean, 2007; Alle et al., 2009; Carter and Bean, 2009; Sengupta et al., 2010; Seutin and Engel, 2010; Carter and Bean, 2011). For instance, the expression of specific subtypes of potassium channels (Kv3 and calcium-activated BK in particular) seems to be necessary to produce sub-millisecond APs (Rudy and McBain, 2001; Lien and Jonas, 2003; Bean, 2007; Alle et al., 2011; Kaczmarek and Zhang, 2017; Hunsberger and Mynlieff, 2020). While there is clear experimental evidence that the properties of sodium and potassium channels shape the AP, the influence of morphological parameters (axon to soma distance, dendritic morphology) on AP properties has been much less investigated. To date, only a few experimental studies and mostly theoretical studies have suggested that differences in morphology may also play a role in the variation of AP duration across neuronal types (Vetter et al., 2001; Kuba et al., 2005; Brette, 2013; Eyal et al., 2014; Gulledge and Bravo, 2016; Kole and Brette, 2018; Goaillard et al., 2020). In addition, the influence of cell-to-cell variations in morphology on AP properties in a same neuronal type has never been investigated experimentally.

In SNc DA neurons, the axon most often arises from an axon-bearing dendrite (ABD) at distances as far as 200 μm from the soma (Hausser et al., 1995; Gentet and Williams, 2007; Meza et al., 2018; Moubarak et al., 2019). The AP, initiated at the axon initial segment, is faithfully back-propagated by the progressive recruitment of somato-dendritic sodium channels (Hausser et al., 1995; Gentet and Williams, 2007; Seutin and Engel, 2010; Moubarak et al., 2019). In this context, we wondered whether the shape of the AP recorded at the soma, its back-propagation efficiency and its modulation by synaptic inputs were influenced by the specific morphologies of the ABD and the non-axon bearing dendrites (nABDs). Using somatic recordings from 38 fully reconstructed neurons, we found that AP HW could be predicted from a linear combination of the respective complexities of the ABD and nABDs (negatively and positively correlated, respectively). The opposite influence of ABD and nABDs was confirmed by selective dendrotomy experiments. Using realistic multi-compartment Hodgkin-Huxley modeling, we then demonstrated that dendritic topology predicts AP HW only when the ABD contains more sodium and/or calcium channels than the nABDs, and that faithful AP back-propagation in response to barrages of EPSCs is also influenced by ABD complexity. In line with previous findings (Moubarak et al., 2019), these results demonstrate that dendritic morphology plays a major role in defining the electrophysiological properties of SNc DA neurons, such that specific cell-to-cell variations in AP dynamics appear to result mainly from variations in dendritic topology.

## MATERIAL AND METHODS

The dataset of neurons presented in this article is the same that was used in a recent publication (6). Additional electrophysiological and morphological parameters, not analyzed for the first article, were extracted from this dataset to perform the current study.

### Acute midbrain slice preparation

Acute slices were prepared from P19-P21 (mean=P20) Wistar rats of either sex (n=19). All experiments were performed according to the European (Council Directive 86/609/EEC) and institutional guidelines for the care and use of laboratory animals (French National Research Council). Rats were anesthetized with isoflurane (CSP) in an oxygenated chamber (TEM SEGA) and decapitated. The brain was immersed briefly in oxygenated ice-cold low-calcium artificial cerebrospinal fluid (aCSF) containing the following (in mM): 125 NaCl, 25 NaHCO_3_, 2.5 KCl, 1.25 NaH_2_PO_4_, 0.5 CaCl_2_, 4 MgCl_2_, 25 D-glucose, or: 87 NaCl, 25 NaHCO_3_, 2.5 KCl, 1.25 NaH_2_PO_4_, 0.5 CaCl_2_, 7 MgCl_2_, 10 D-glucose, 75 sucrose; pH 7.4, oxygenated with 95% O_2_/5% CO_2_ gas. The cortices were removed and then coronal midbrain slices (250 or 300 μm) were cut on a vibratome (Leica VT 1200 or 1200S) in oxygenated ice-cold low calcium aCSF. Following 20-30 min incubation in 32°C oxygenated low calcium aCSF the slices were incubated for at least 30 minutes in oxygenated aCSF (125NaCl, 25 NaHCO_3_, 2.5 KCl, 1.25 NaH_2_PO_4_, 2 CaCl_2_, 2 MgCl_2_ and 25 glucose, pH 7.4, oxygenated with 95% O_2_ 5% CO_2_ gas) at room temperature prior to electrophysiological recordings.

### Drugs

Picrotoxin (100μM, Sigma Aldrich) and kynurenate (2mM, Sigma Aldrich) were bath-applied via continuous perfusion in aCSF to block inhibitory and excitatory synaptic activity, respectively.

### Electrophysiology recordings and analysis

All recordings (45 cells from 19 rats) were performed on midbrain slices continuously superfused with oxygenated aCSF. Patch pipettes (1.8-3 MOhm) were pulled from borosilicate glass (GC150TF-10, Havard Apparatus) on a DMZ Universal Puller (Zeitz Instruments). The patch solution contained in mM: 20KCl, 10 HEPES, 10 EGTA, 2 MgCl2, 2 Na-ATP and 120 K-gluconate, pH 7.4, 290-300 mOsm. Neurobiotin (0.05%; Vector Labs) was included in the intracellular solution to allow morphological reconstruction and identification of dopaminergic neurons using post-hoc tyrosine-hydroxylase immunolabeling (Amendola et al., 2012; Moubarak et al., 2019). Whole-cell recordings were made from SNc DA neurons visualized using infrared differential interference contrast videomicroscopy (QImaging Retiga camera; Olympus BX51WI microscope) and identified as previously described (Amendola et al., 2012; Moubarak et al., 2019). Whole-cell current-clamp recordings with a series resistance <10MOhm were included in the study. Capacitive currents and liquid junction potential (+13.2mV) were compensated online and offset potentials were measured after removing the pipette from the neuron. Bridge balance (100%, 10μs) was used to compensate series resistance. Recordings with offset values above 1mV were discarded from the analysis. Recordings were acquired at 50kHz and were filtered with a low-pass filter (Bessel characteristic between 2.9 and 5kHz cutoff frequency). Action potentials (APs) generated during a 40s period of spontaneous activity were averaged and the AP threshold, AP amplitude and the duration of the AP at half of the maximal height of the AP (AP half-width, AP HW) were measured. AP threshold was measured on the d^2^v/dt^2^ *vs* V phase plane plot.

### Laser photo-ablation of dendrites

For dendrotomy experiments, Alexa 594 (4%, Invitrogen) and neurobiotin (0.05%, Vector Labs) were included in the patch solution, and confocal live imaging was performed on an LSM 710 Zeiss confocal microscope (Carl Zeiss AG, Germany), with the images being captured using Zeiss ZEN software. Recordings and live imaging were started 10-15 minutes after obtaining the whole-cell configuration in order to allow substantial diffusion of the dye into the dendritic tree. To prevent fatal damage of the neuron, a region of interest (ROI) was then chosen on a secondary or tertiary dendrite, at least 50 μm from the soma. A z-stack image of the neuron comprising the ROI and the soma was taken before starting the laser photo-ablation procedure. The ROI was then illuminated for 10-15s with medium power using a 543 nm laser. After a 1-5 minutes recovery period, images of the ROI were taken to assess ablation success. Dendrotomy is characterized by membrane swelling in the ROI and loss of upstream and downstream fluorescence (Go et al., 2016; Rama et al., 2017). The illumination procedure was repeated as many times as necessary until ablation was achieved. After dendrotomy was obtained, a z-stack image of the neuron comprising the ROI and the soma was taken, and compared with the image taken before laser illumination. APs were induced by incremental step current injections and recorded before and after each laser illumination.

### Electrophysiology data acquisition and analysis

Data were acquired with a HEKA EPC 10/USB patch-clamp amplifier (HEKA electronics) and patchmaster software (HEKA electronics) or a Multiclamp 700B (Molecular Devices, Palo Alto, CA). Analysis was conducted using FitMaster v2×30 (Heka Elektronik) and Clampfit (Molecular Devices).

### Immunohistochemistry and morphological reconstruction

Acute slices containing Neurobiotin-filled cells were fixed 30 min in 4% paraformaldehyde at 4°C and immunolabelled with anti-tyrosine hydroxylase (chicken polyclonal, Abcam, 1:1000) and anti-AnkyrinG (mouse monoclonal IgG2b, NeuroMab, 1:250) antibodies. Goat anti-mouse IgG2b Alexa Fluor 488 (Invitrogen; 1:1000; 2μg/ml) and goat anti-chicken Alexa Fluor 633 (Invitrogen; 1:3000; 1.66ng/ml) were used to reveal tyrosine hydroxylase and AnkyrinG stainings, respectively. Streptavidin Alexa Fluor 594 (Invitrogen; 1:12000; 1.66ng/ml) was used to reveal neurobiotin labelling. Slices were mounted in Faramount mounting medium (Dako). Immunolabellings were viewed on an LSM 780 Zeiss (Carl Zeiss AG, Germany) and images were captured using Zeiss ZEN software. Images were analyzed with Fiji/ImageJ software (Schindelin et al., 2012; Schneider et al., 2012; Rueden et al., 2017) and in particular neurons were reconstructed using the Simple Neurite Tracer plugin (Longair et al., 2011). The axon was identified using ankyrinG labelling, which allowed us to discriminate between ABD and nABDs. All dendritic lengths were extracted directly from the paths traced through the stack images of the neurons, this measure corresponding to the sum of the lengths of all dendritic segments arising from a given primary dendrite (ABD or nABD). To assess dendritic complexity, we extracted from the reconstructions the number of dendritic segments (branches) found on the ABD and nABDs. The average length and number of segments per nABD were obtained by dividing the total for these parameters by the number of primary nABD dendrites for each neuron. Soma volumes were estimated by using the “fill out path” method on Simple Neurite Tracer. For the diameters of the primary dendrites, intermediate ABD dendrites, axon and AIS, the fluorescence histograms of branch sections were obtained and fitted with a Gaussian curve model. Diameters were then estimated as 3 times the standard deviation (3*SD) of the Gaussian distribution.

### Multi-compartment modelling

Simulations were performed using NEURON 7.5 software (Hines and Carnevale, 1997, 2001; Carnevale and Hines, 2005), as described in a previous publication (Moubarak et al., 2019). Neuronal morphologies correspond to the realistic morphologies from the 37 fully reconstructed dopaminergic neurons.

For each compartment, membrane voltage was obtained as the time integral of a first-order differential equation:

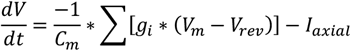

where *V_m_* is the membrane potential, *C_m_* the membrane capacitance, *g_i_* are ionic conductances and *V_rev_* their respective reversal potentials. The axial flow of current (I axial) between adjacent compartments is calculated by the NEURON simulation package (Hines and Carnevale, 1997). Cytoplasmic resistivity, specific membrane capacitance and specific membrane resistance were set to 150 Ohm.cm, 0.75 μF/cm^2^, and 100,000 Ohm*cm^2^, respectively, with the reversal potential for the leak conductance set at −50 mV. Active conductances followed activation-inactivation Hodgkin-Huxley kinetics (**Table 1**).

**TABLE 1.**
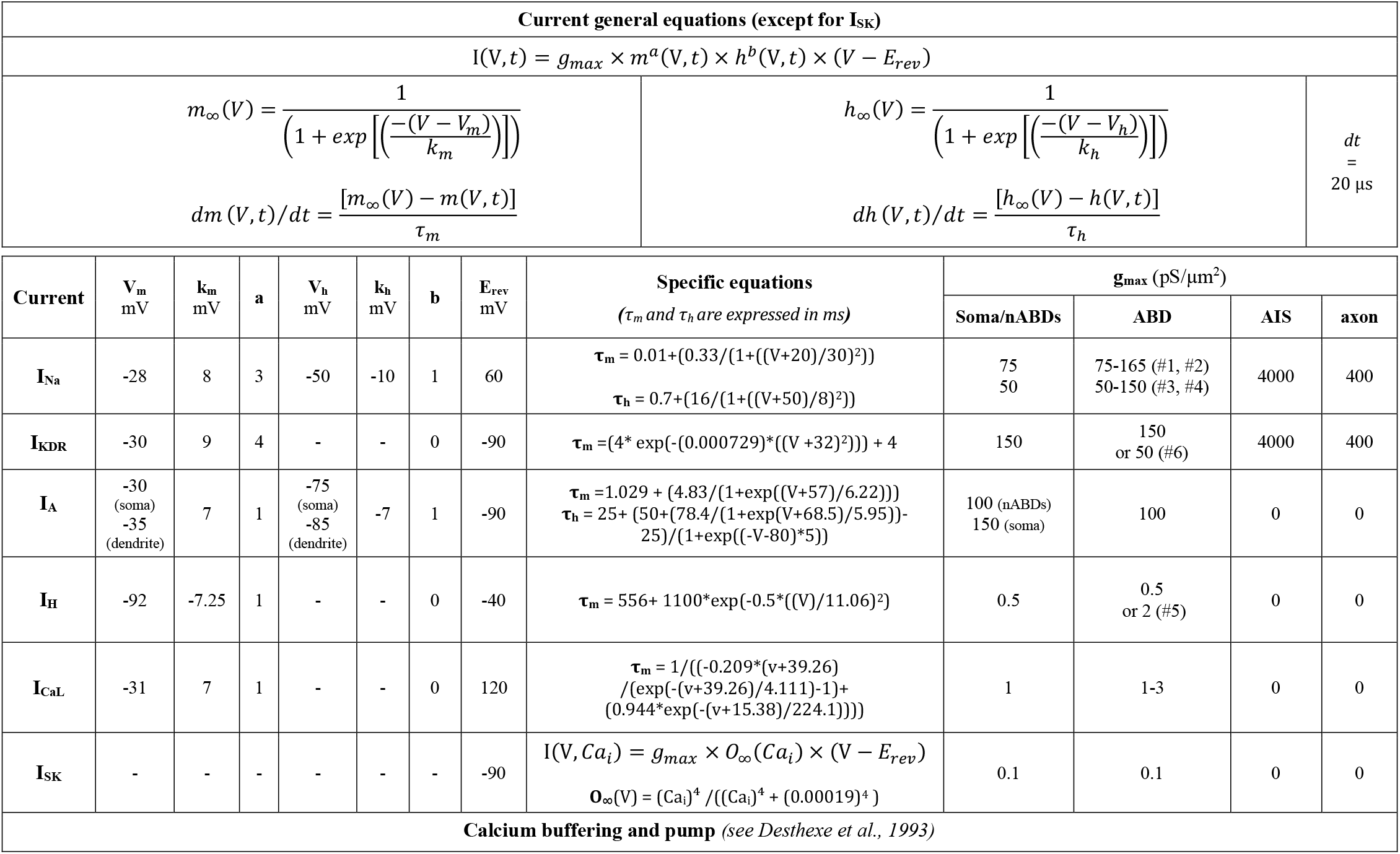
Equations governing the voltage dependence and kinetics of currents in the model. The 6 versions of the model implementing different values of g_Na_, g_KDR_, g_H_ or g_CaL_ in the ABD are indicated (#) with the corresponding conductance values or ranges.

Parameters for I_A_, I_CaL_, I_KCa_ and I_H_ were based on previous published values for SNc DA neurons (Wilson and Callaway, 2000; Amendola et al., 2012; Engel and Seutin, 2015). Fast sodium and potassium currents were derived from Migliore and Schild models (Schild et al., 1993; Migliore et al., 2008), respectively. The SK current is solely activated by an increase in calcium concentration. Therefore, intracellular calcium uptake was modeled as a simple decaying model according to (Destexhe et al., 1993). Conductance values were set according to our own measurements or published values (see **Table 1**). Consistent with the literature (Zhou et al., 1998; Kole et al., 2008; Hu et al., 2009; Gonzalez-Cabrera et al., 2017), g_Na_ and g_KDR_ densities are higher in the AIS than in the rest of the neuron so that AP always initiates in the AIS. According to Gentet and Williams (2007), I_A_ density and inactivation kinetics were higher and depolarized, respectively, in the soma compared to the dendritic arbor. Initializing potential was set at −70 mV and analysis was performed after pacemaking frequency reached a steady-state (8 spikes). Each simulation run had a 6000 ms duration with a dt of 0.01 ms. All dendritic compartments and the axon-start compartment contained all currents whereas AIS and axon only contained fast sodium and potassium currents (g_Na_, g_KDR_). Unless otherwise stated, all currents but the fast sodium and potassium currents had fixed and homogeneous conductance values in the dendrites and the axon-start compartment.

For the realistic morphology models, exact dendrite lengths, soma volume, and diameters of primary dendrites, aDs, axon and AIS were used (see (Moubarak et al., 2019) for details). The specific branching patterns of each neuron (topology) were also respected. In order to be consistent with the NEURON software constraints, soma volume was implemented by computing the equivalent cylinder corresponding to the volume measured using “fill out path” method on Simple Neurite Tracer. Axonal diameter was considered constant and set to 0.7μm while the diameters of non-primary dendrites were approximated by a regular tapering to reach a final diameter of 0.5μm.

Firing frequency and AP analysis (amplitude, first and second derivative of APs) were computed online by handmade routines directly written in NEURON hoc language. For the dendrotomy simulation, dendritic cut was modeled by decreasing the diameter of the cut branch to nearly zero (1 pm). EPSPs were modeled as alpha synapses with a rise time of 0.1 ms, a decay time of 1 ms and a reversal potential of 0 mV (Gentet and Williams, 2007). The frequency of barrages of simulated EPSPs were designed according to Gentet and Williams as the convergence of five neurons firing irregularly at a mean frequency of 20 Hz, following a negative binomial distribution. All computing files are available at model DB database (entry #245427).

### Experimental design and statistical analysis

Statistical analysis (performed according to data distribution) included: linear regression, multiple linear regression, unpaired t-test, Mann Whitney, paired t-test and one-way ANOVA, with a p value <0.05 being considered statistically significant. Distribution of data was first tested for normality with the Shapiro–Wilk test. Then, the difference between means of two samples was accordingly computed using t-tests or Wilcoxon signed rank tests. For comparison between three groups, we used a one-way parametric ANOVA followed by post hoc t-tests with Bonferroni correction for multiple comparisons. Linear regressions were obtained with the Pearson test. Multiple regressions were computed as a linear model such as HW= a*ABD +b* ABD + error. P values for regressions were corrected for multiple comparisons with a false discovery rate procedure (Benjamini and Hochberg, 1995). Predictive values of the multiple linear models were tested by multiple repeated cross-validation (R, caret package). Briefly, data were randomly partitioned into train and test data (80%/20%). Repeated (20 times) 10-fold cross-validation was computed on train data and the model was tested against the test data to calculate the r. This procedure was repeated 100 times in order to calculate the mean and confidence interval (95%) for r and the model coefficients. Unless otherwise stated, statistical data are given as mean ± standard deviation and n indicates the number of recorded neurons. Statistical tests were computed by using Sigmaplot 11.0 software (Systat Software Inc., San Jose, CA, USA) and R.

### Figure preparation

Figures were prepared using Sigma Plot, Prism 6, Adobe Photoshop and Adobe Illustrator (CS5-CS6, Adobe Systems Inc., San Jose, CA, U.S.A.), and ImageJ (Schindelin et al., 2012; Schneider et al., 2012; Rueden et al., 2017), with brightness and contrast adjustments performed consistently across the images to enhance clarity.

## RESULTS

To explore the link between AP properties and morphology in SNc DA neurons, we performed somatic patch-clamp recordings combined with post-hoc reconstructions of neuronal morphology based on neurobiotin fills of the recorded neurons (n=38). AnkyrinG immunostaining was used to identify the axon (Meza et al., 2018; Moubarak et al., 2019), and the morphological properties of the ABD and nABDs were analyzed (**Figure 1, Figure 2**). In SNc DA neurons, the AP is initiated at the AIS and back-propagates sequentially through ***i)*** the ABD stem, ***ii)*** the soma and ***iii)*** the nABDs 25. This back-propagation relies on somato-dendritic sodium channels (Hausser et al., 1995; Seutin and Engel, 2010; Moubarak et al., 2019), which also participate in the genesis of spontaneous pacemaking activity (Tucker et al., 2012; Jang et al., 2014; Moubarak et al., 2019). We therefore wondered whether the shape of the AP recorded at the soma could be influenced by the morphological properties of the dendritic arbor (**Figure 1**).

**Figure 1.**
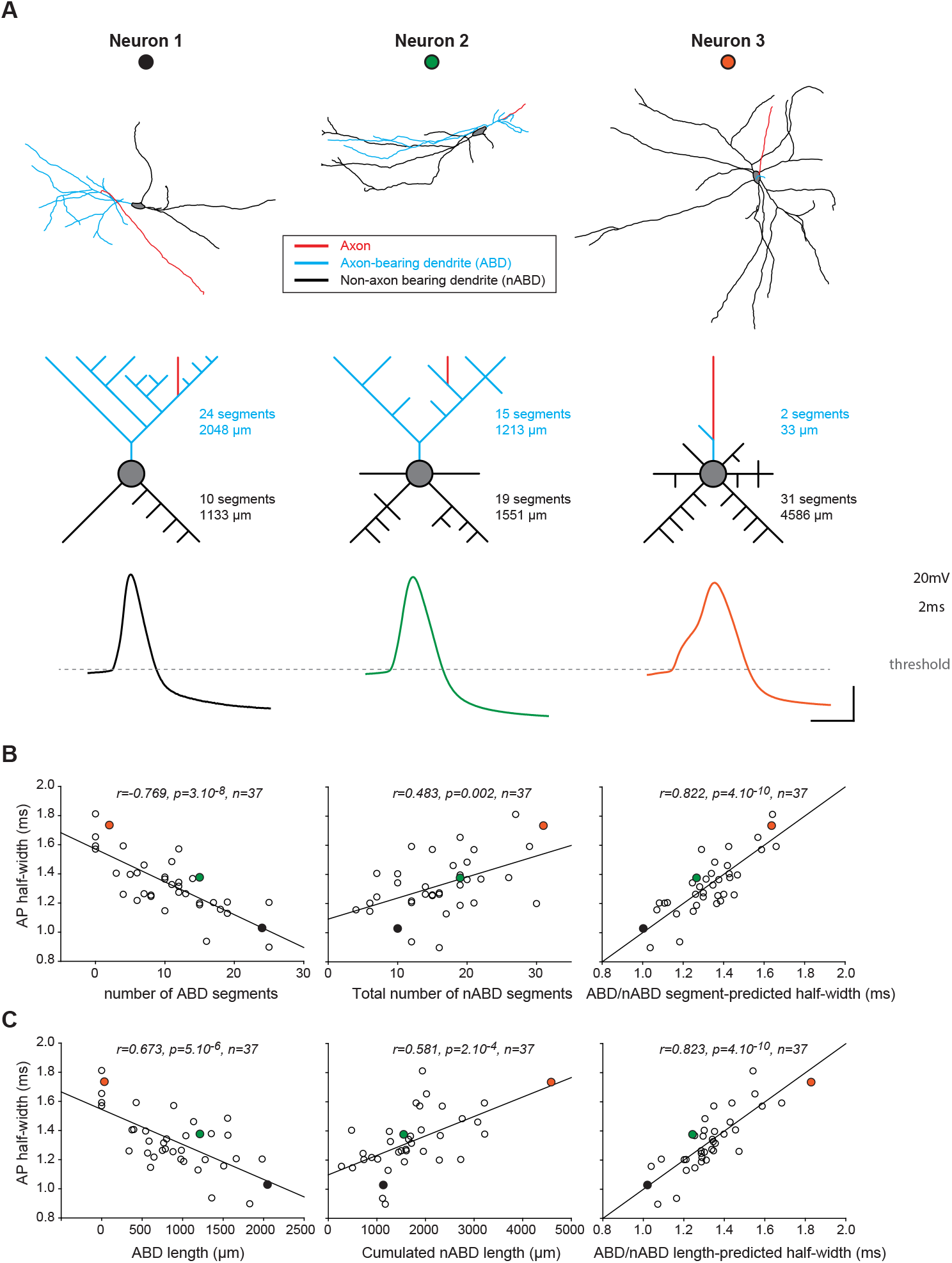
Dendritic topology predicts somatic action potential half-width in SNc DA neurons. **A**, **top row**, skeleton pictures of 3 fully reconstructed SNc DA neurons. **Middle row**, schematic representations of the topology corresponding to the same 3 neurons. The number of dendritic segments comprising the ABD (blue) and the nABDs (black) as well as their respective lengths are indicated. Bottom row, somatic current-clamp recordings of the AP obtained in the same 3 neurons. Note that the action potential is getting slower when the complexities of the ABD and nABDs are decreasing and increasing, respectively. **B**, **left**, scatter plot showing the negative correlation between ABD complexity and AP half-width. **Center**, scatter plot showing the positive correlation between nABD complexity and AP half-width. **Right**, scatter plot showing the correlation obtained using a linear combination of ABD and nABD complexities (multiple linear correlation). **C**, **left**, scatter plot showing the negative correlation between ABD length and AP half-width. **Center**, scatter plot showing the positive correlation between nABD length and AP half-width. **Right**, scatter plot showing the correlation obtained using a linear combination of ABD and nABD lengths (multiple linear correlation). The black, green and orange dots correspond to the specific values obtained for the 3 neurons presented in panel **A**. Scale bars: **A**, vertical 20mV, horizontal 2ms. The horizontal dotted line indicates action potential threshold.

**Figure 2.**
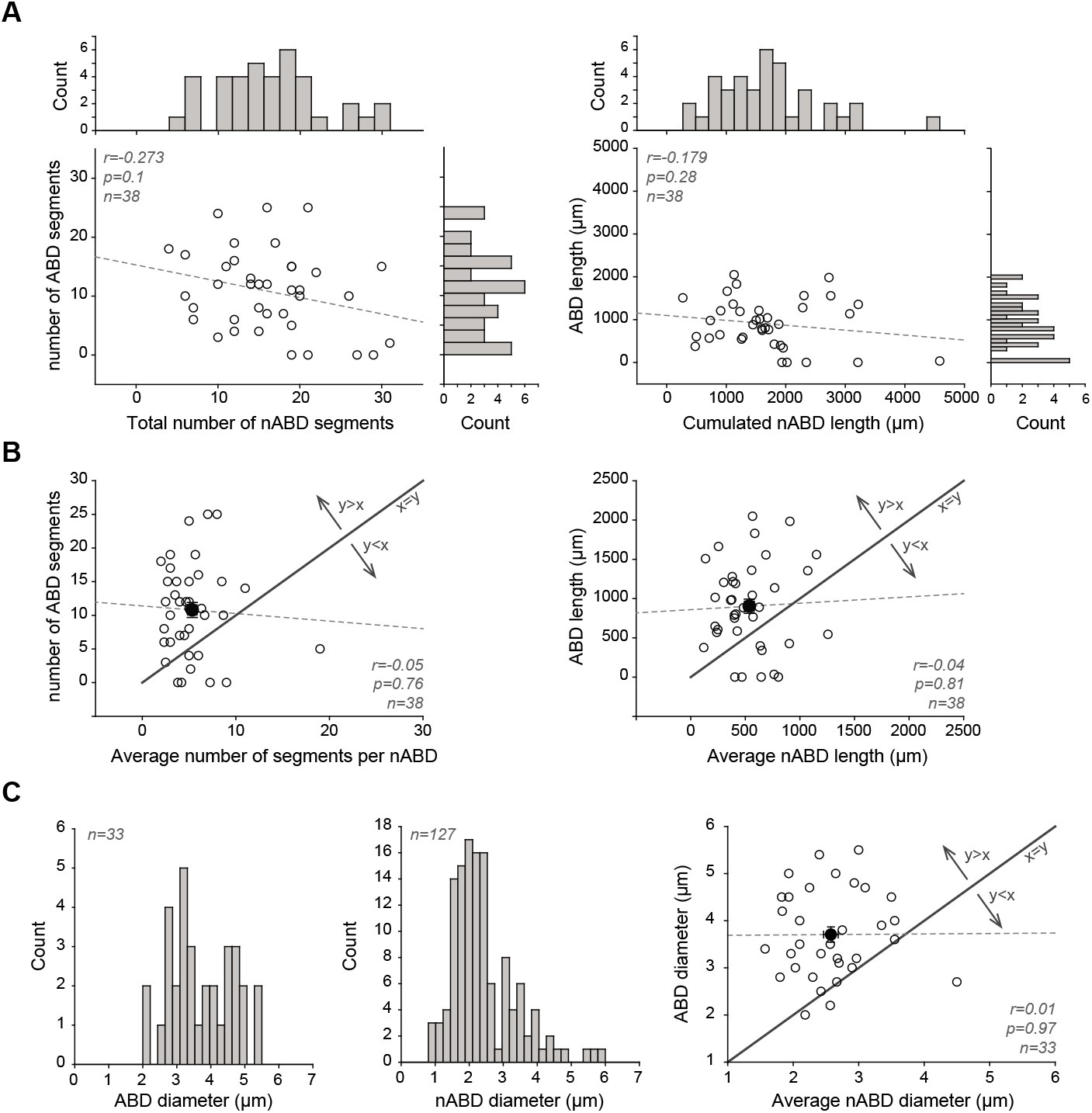
Detailed description and comparison of ABD and nABD morphological properties. **A**, **left**, scatter plot representing the lack of relationship between the total complexity of the nABDs and the complexity of the ABD. Histograms showing the distribution of values for the number of segments of the ABD and nABDs are displayed on the right and top of the scatter plot, respectively. **Right**, scatter plot showing the lack of relationship between the cumulated length of the nABDs and the length of the ABD. Histograms showing the distribution of values for the length of the ABD and nABDs are displayed on the right and top of the scatter plot, respectively. **B**, **left**, scatter plot representing the complexity of the ABD versus the average complexity of a nABD. The black dot corresponds to the average measured in 38 neurons. The plain line corresponds to the identity line. Please note that the number of segments per nABD is not correlated with the number of segments per ABD, and that the number of segments per ABD is on average larger than the number of segments per nABD. **Right**, scatter plot representing the length of the ABD versus the average length of a nABD. The black dot corresponds to the average measured in 38 neurons. The plain line corresponds to the identity line. Please note that the average length of a nABD is not correlated with the length of the ABD, and that the length of the ABD is on average larger than the average length of a nABD. **C**, **left and middle**, histograms showing the distribution of values for the diameters of the ABD and the nABDs measured in 33 and 37 neurons, respectively. Right, scatter plot representing the diameter of the ABD versus the average diameter of a nABD. Please note that the average diameter of a nABD is not correlated with the diameter of the ABD, and that the diameter of the ABD is on average larger than the average diameter of a nABD.

We focused in particular on the overall length of the ABD and nABDs as well as the number of dendritic segments comprising them (**Figure 1A, Figure 2**). This analysis was performed on the 34 neurons where the axon arose from an ABD (34/38~90%). The ABD was significantly longer (1008±495μm *vs* 532±283μm; t=5.073, p<0.001, n=34, paired t test) and more complex (containing more dendritic segments; 12.1±6.1 *vs* 5.2±3.2; t= 5.743, p<0.001, n=34, paired t test) than any nABD (**Figure 2**). On average the starting diameter was also found to be significantly larger for the ABD than for the nABDs (3.7±0.9μm *vs* 2.8±0.6μm; t= 5.774, p<0.001, n=34, paired t test). Notwithstanding these differences in the average morphology of the ABD and nABDs, the relative contribution of the ABD and nABDs to the overall dendritic tree was extremely variable from neuron to neuron (**Figure 1A, Figure 2**). As previously described (Amendola et al., 2012; Dufour et al., 2014; Moubarak et al., 2019), the duration of the AP recorded at the soma also displayed considerable cell-to-cell variations, ranging from 0.9 to 1.81 ms (1.33±0.20ms, n=37). Intriguingly, we found that AP HW was significantly correlated, albeit in opposite ways, with the complexity and the length of the ABD and nABDs (**Figure 1B, C**). Specifically, the number of dendritic segments in the ABD and its length were negatively correlated with AP HW while the same parameters for the nABDs were positively correlated with AP HW (**Figure 1B, C**). The morphological properties of the ABD and nABDs were independent from each other (**Figure 2**) and we combined them to perform multiple linear regressions against AP HW (**Figure 1B, C**). Using ABD/nABD complexity or length, AP HW could be predicted with a Pearson correlation coefficient > 0.82. Multiple linear models obtained by cross-validation (see methods) predicted a correlation with similar r values (r=0.75, CI_95_=0.63-0.86). While AP HW is a “standard” parameter used to describe AP shape, it is merely a phenomenological measurement that captures the combined variations in rising and decaying rates of the AP (HW=-1.92*(rise slope)+19.4*(decay slope); all terms significant p<0.001, adjusted r = 0.898, p<2.10^-16^, n=51, multiple linear regression), which are more closely related to the density of inward and outward currents underlying the AP (Bean, 2007). Consistent with this, both rising and decay rates were also significantly correlated with ABD complexity (r=0.41, p=0.0011 and r=-0.54, p=0.0005, respectively). Thus, although ion channel properties and densities vary from cell to cell (Liss et al., 2001; Seutin and Engel, 2010; Amendola et al., 2012; Moubarak et al., 2019; Haddjeri-Hopkins et al., 2021) and have been suggested to have a significant influence on AP back-propagation (Moubarak et al., 2019), knowing the specific dendritic topology of an SNc DA neuron (complexity of the ABD and nABDs) is sufficient to predict 68% of the variance in duration of the AP at the soma, suggesting that dendritic topology largely constrains AP duration. None of the other parameters defining AP shape (amplitude, threshold, AfterHyperPolarisation voltage) shared a significant correlation with morphological measurements.

In order to investigate the potential biophysical underpinnings of this unexpected relationship, we then used a multi-compartment Hodgkin-Huxley model based on the precise morphological measurements obtained from the 37 neurons where electrophysiological recordings were performed (for details see (Moubarak et al., 2019)). The opposite correlations shared by the ABD or nABDs with AP HW suggest contrasting influences of these dendritic branches on AP shape. While the ABD can be viewed as an accelerator of the AP (complexity/length negatively correlated with AP HW), the nABDs seem to slow down the AP (complexity/length positively correlated with AP HW). Another way of seeing it is that the ABD has an “active” influence on the shape of the somatic AP while the nABDs mainly acts as a “passive sink” (Brette, 2013; Kole and Brette, 2018). The sodium and L-type calcium channels are the main depolarizing conductances defining the upstroke of the AP in SNc DA neurons (Bean, 2007; Puopolo et al., 2007; Seutin and Engel, 2010; Philippart et al., 2016). We therefore tested whether the opposite correlations between ABD or nABDs and AP HW could be explained by a heterogeneous density of these conductances in the dendritic tree. The average density of sodium conductance (g_Na_) in the dendrites of rat SNc DA neurons, measured in a previous study (Moubarak et al., 2019), has been estimated to be ~75 pS/μm^2^, while the dendritic density of L-type calcium current (g_Ca_) is unknown. We linearly and independently increased g_Na_ and g_Ca_ in the ABD up to ~3-fold while keeping their density constant in the soma and nABDs, starting with a baseline g_Na_ of either 75 or 50 pS/μm^2^ and a baseline g_Ca_ of 1 pS/μm^2^ (**Figure 3, Figure 4**). Using this approach, a database of 231 models for each of the 37 reconstructed neurons was created (**Figure 4A**). In all conditions, these models of SNc DA neurons still generated a pacemaking pattern of activity in the expected range of frequencies (**Figure 3B**). We then analyzed whether the correlations between ABD or nABD complexity and AP HW observed in the real neurons were reproduced in the model and how they were influenced by increasing g_Na_, g_Ca_ or both, in the ABD compared to the nABDs (**Figure 3C, Figure 4A**). While the positive correlation between nABD complexity and AP HW was fairly insensitive to changes in ABD g_Na_ and g_Ca_, the negative correlation between ABD complexity and AP HW was present only when ABD g_Na_ (or/and g_Ca_) was increased (**Figure 3C**). In fact, simultaneously increasing both conductances very much improved the power of this correlation, such that the multiple linear regression based on ABD and nABD complexities was significant only in this case (**Figure 3C**, **Figure 4A**), with a Pearson correlation factor close to the one observed in real neurons (r=0.760, p=7.10^-8^, n=36 in the model compared to r=0.822, p=4.10^-10^, n=37 in the real neurons). Multiple linear models obtained by cross-validation (see methods) predicted a correlation with similar r values (r=0.60, CI_95_=0.52-0.70). Consistent with the experimental results reported before (**Figure 1C**), similar results were obtained when using the length of the ABD and nABDs instead of their complexity (data not shown). Most interestingly, in the heterogeneous condition (g_Na/_g_Ca_ increased in the ABD) the half-width of the AP produced by the model neurons predicted the half-width of the AP recorded in the real neurons (r=0.718, p=1.10^-6^, n=35, **Figure 3C**). Since other conductances are involved in defining AP duration (such as the delayed rectifier potassium current) or have been suggested to display a higher conductance density in the ABD (such as I_H_; see (Engel and Seutin, 2015)), we tested whether increasing ABD excitability by the manipulation of these conductances (decreasing g_K_ or increasing g_H_ in the ABD) had a similar effect on the observed correlations. While the positive correlation between nABD complexity and AP HW was present in all conditions, the negative correlation between ABD complexity and AP HW was not reproduced in models where g_H_ or g_K_ was increased or decreased, respectively, in the ABD (**Figure 4B, C**). Thus, the correlation between dendritic topology and AP duration observed in real neurons can only be reproduced in a realistic model if g_Na_ and g_Ca_ specifically are larger (by ~2-fold) in the ABD compared to the nABDs. Moreover, this condition of heterogeneous dendritic excitability allows the model to predict the duration of the somatic AP recorded in the real neurons. Since modifications in dendritic biophysical properties (increased g_Na_/g_Ca_) were identical for all model neurons, this result demonstrates that cell-to-cell variations in dendritic topology are sufficient to explain most of the variation in somatic AP shape observed in real neurons.

**Figure 3.**
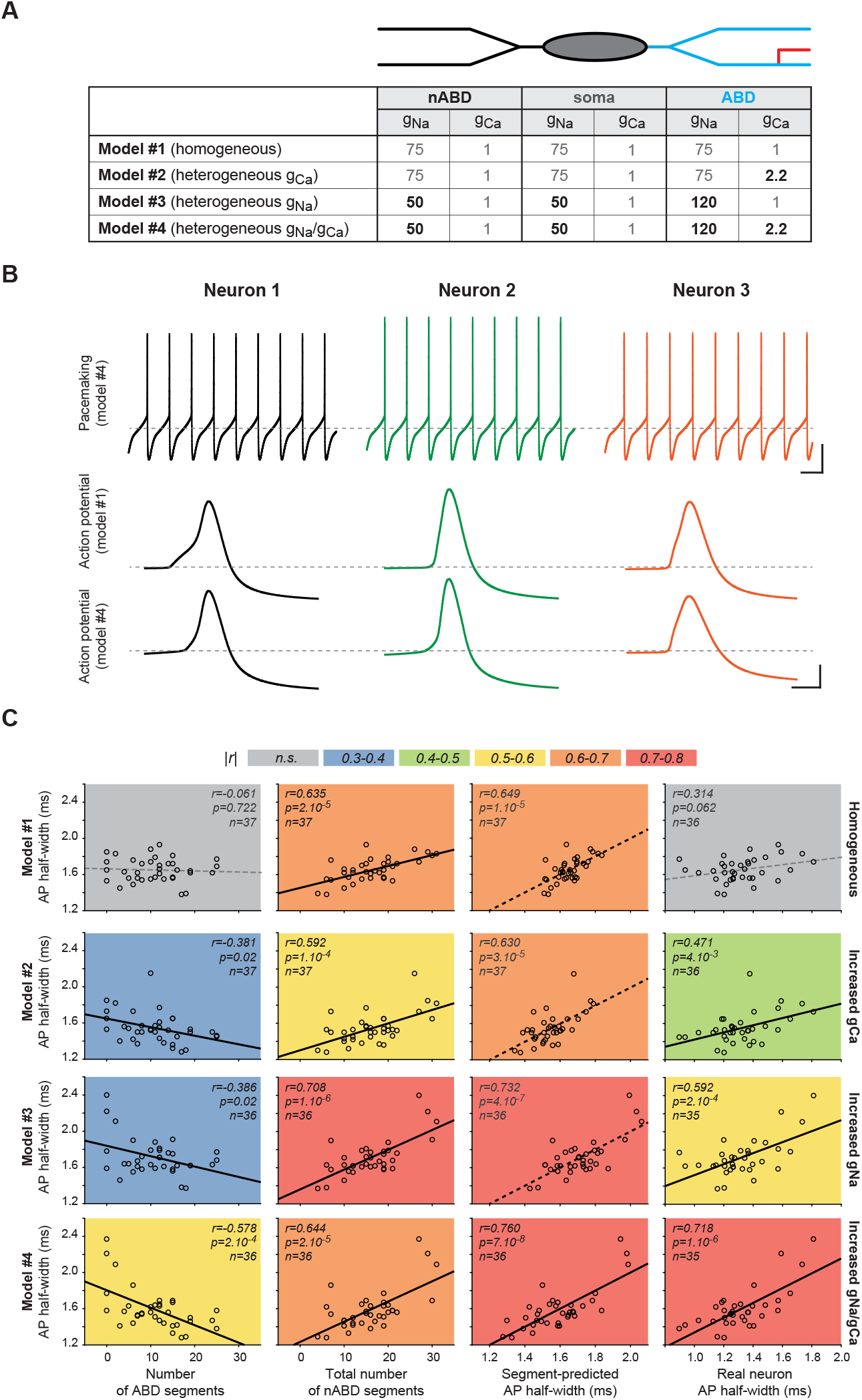
A realistic morphology model of SNc DA neurons reproduces the correlations between dendritic topology and AP half-width and predicts the recorded AP half-width. **A**, table describing the specific values of sodium (g_Na_) and calcium conductance (g_Ca_) implemented in the nABDs, soma and ABD in the 4 versions of the model presented in the rest of the figure. **B**, **top row**, example traces showing the voltage waveforms of spontaneous activity produced by the model #4 for the models of the 3 neurons presented in **Figure 1**. **Middle and bottom rows**, AP waveforms obtained for the same neurons for the models #1 and #4, respectively. **C**, scatter plots representing the relationships between ABD complexity (first left), nABD complexity (second left), a linear combination of ABD and nABD complexity (second right) or the AP half-width measured in real neurons (first right) and the AP half-width measured in the realistic morphology models. The relationships are shown for the 4 versions of the model (top to bottom) presented in panel **A**. Non-significant correlations are indicated by gray dotted lines. Scatter plots are color-coded, depending on the value of the correlation coefficient. Colored dotted lines (multiple linear correlations) indicate that only one variable (nABD complexity) significantly participates in the correlation. Scale bars: **B**, top traces, vertical 20mV, horizontal 200ms; middle and bottom traces, vertical 20mV, horizontal 2ms. Horizontal gray dotted lines indicate −60mV (top traces) and action potential threshold (middle and bottom traces).

**Figure 4.**
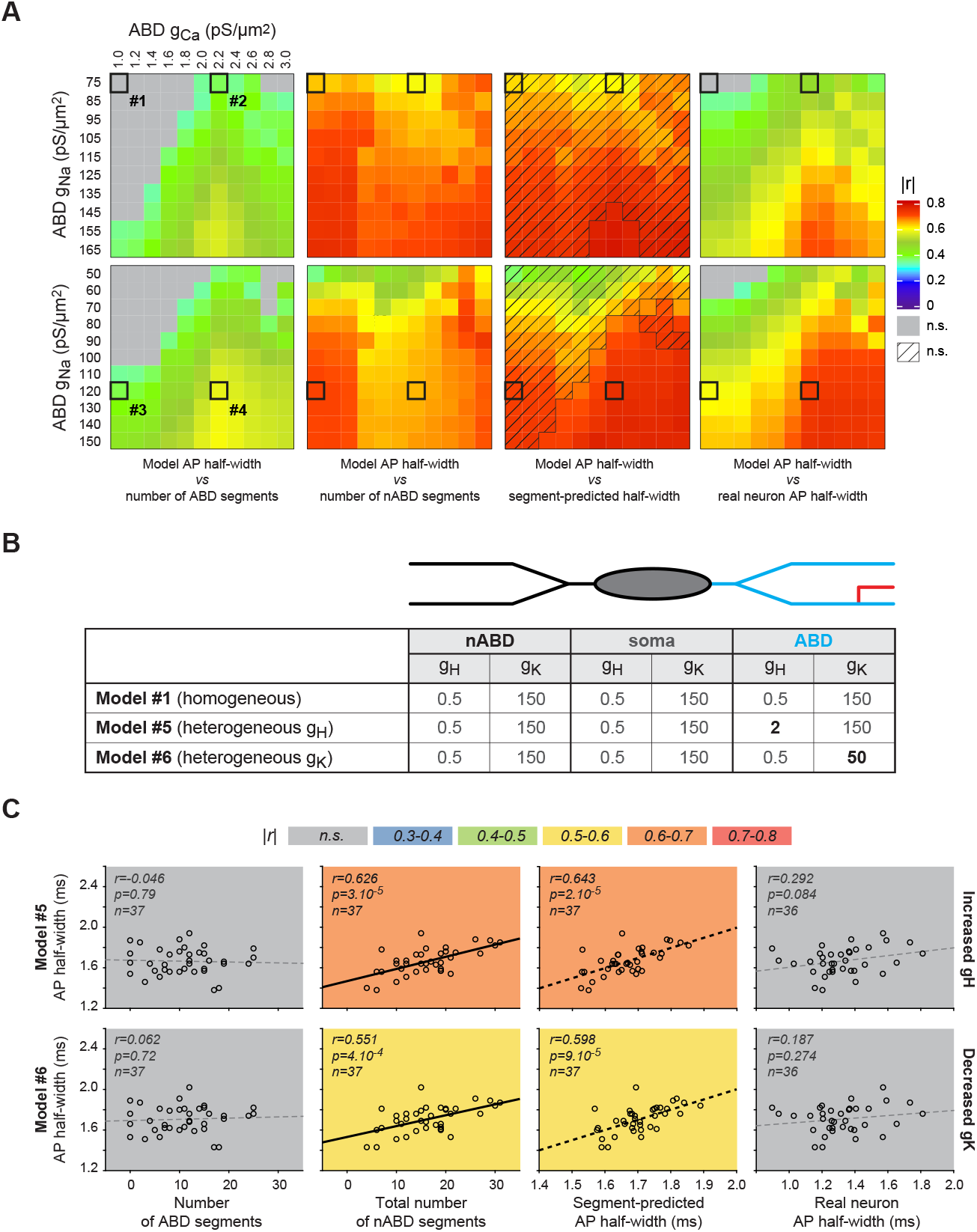
Exploring the biophysical conditions necessary to reproduce the correlation between dendritic topology and AP half-width. **A**, heatmaps representing the evolution of the correlation factor r for the relationships presented in **Figure 2C** when independently and linearly varying g_Ca_ in the ABD from 1 to 3 pS/μm^2^ (top and bottom heatmaps) and g_Na_ in the ABD from 75 to 165 pS/μm^2^ (top) or from 50 to 150 pS/μm^2^ (bottom). nABD conductance values were kept fixed at 1 and 75 pS/μm^2^ (top) or 1 and 50 pS/μm^2^ (bottom) for g_Ca_ and g_Na_, respectively. The 4 versions of the model presented in Figure 2 are highlighted by a dark surrounding. The shaded area in the heatmap corresponding to the multiple linear regression (second right) indicates that only one variable (nABD complexity) significantly participates in the correlation. **B**, table describing the specific values of I_H_ (g_H_) and delayed rectifier potassium conductance (g_K_) implemented in the nABDs, soma and ABD in the 2 versions of the model presented in panel **C**. **C**, scatter plots representing the relationships between ABD complexity (first left), nABD complexity (second left), a linear combination of ABD and nABD complexity (second right) or the AP half-width measured in real neurons (first right) and the AP half-width measured in the realistic morphology models. The relationships are shown for the 2 versions of the model (#5, top; #6, bottom) presented in panel **B**. Non-significant correlations are indicated by gray dotted lines. Scatter plots are color-coded, depending on the value of the correlation coefficient. Colored dotted lines (multiple linear correlations) indicate that only one variable (nABD complexity) significantly participates in the correlation.

So far, the relationships between dendritic topology and AP duration observed in the real neurons and their realistic computer-simulated counterparts are only correlative. However, since our model is purely deterministic, these correlations suggest that increasing the complexity/length of the ABD indeed “speeds up” the AP while increasing the complexity/length of the nABDs “slows it down”. To test the causality of this link, we simulated ABD and nABD dendrotomies and determined their sensitivity to g_Na_/g_Ca_ conductance distribution (**Figure 5**). ABD and nABD dendrotomies were simulated on the same neurons using both the homogeneous (same g_Na_/g_Ca_ in ABD and nABDs, model #1) and heterogeneous (increased g_Na_/g_Ca_ in the ABD, model #4) versions of the model. On the ABD side, dendrotomies of intermediate dendrites (located in between the axon and the soma) or distal dendrites (more distal from the soma than the axon) were tested (see **Figure 5A**): these two manipulations produced similar effects on AP HW and were thus pooled in our statistical analysis. Simulating a dendrotomy of the ABD or nABDs in the homogeneous model had the same effect on AP HW, inducing a small but significant decrease in AP HW (ABD, 100 *vs* 98.14%, t=6.878, p<0.001, n=44, paired t-test; nABD, 100 *vs* 96.26%, z=-5.159, p<0.001, n=37, signed rank test; **Figure 5B, C**). However, in the heterogeneous version of the model, ABD and nABD dendrotomies had opposite effects on AP HW: sectioning the nABD still induced a decrease in AP HW (100 *vs* 94.92%, z=-5.16, p<0.001, n=35, signed rank test; **Figure 5B,C**), while sectioning the ABD significantly increased AP HW (100 *vs* 102.55%, t=6.618, p<0.001, n=41, paired t-test; **Figure 5C**). Thus, an increased g_Na_/g_Ca_ in the ABD confers opposite causal influences of the ABD and nABDs on AP duration, hence corroborating the opposite correlations observed between ABD or nABDs and AP HW in real neurons and in the models (**Figure 1, Figure 3**).

**Figure 5.**
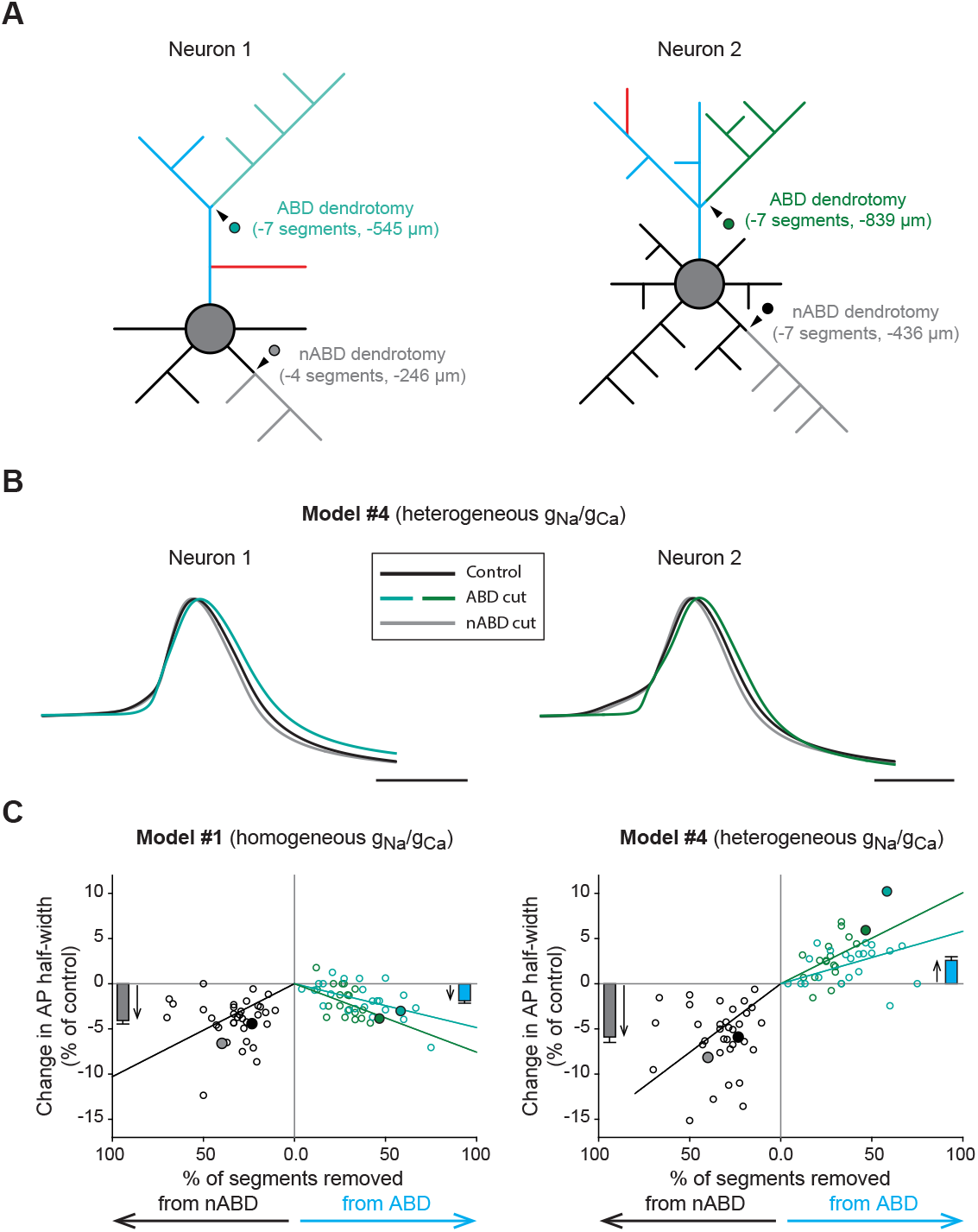
Modeling the effect of ABD and nABD dendrotomies. **A**, schematic representations of the topology of 2 neurons, indicating the location and extent of the dendrotomies simulated on the ABD (green dots) or nABD (gray and dark dots). **B**, voltage waveforms of the AP obtained for the model #4 of neuron 1 (left) and neuron 2 (right) before dendrotomy (black trace), and after ABD (green) or nABD dendrotomy (gray). AP waveforms were scaled and aligned at midamplitude to allow comparison of the half-widths. **C**, scatter plots representing the change in AP half-width following simulated dendrotomy in the model #1 (left) and #4 (right) as a function of the percentage of nABD or ABD segments removed from the model. The average changes in AP half-width obtained for nABD and ABD dendrotomies are indicated as bar plots (mean±SEM) on each scatter on the left and right, respectively. Please note that only the model #4 predicts an opposite influence of nABDs and ABD on AP half-width. Scale bars: **B**, 2ms.

Based on this prediction from the model, we sought to determine whether experimentally reducing dendritic complexity by specific ABD and nABD dendrotomies would have opposite effects on AP HW in real neurons (**Figure 6**). In order to test this hypothesis, neurons were filled with Alexa 594 during patch-clamp recording, and laser illumination (line scan) was used to induce dendrite photo-ablation (**Figure 6A**). The nature of the sectioned dendrite (ABD *vs* nABD) was determined by post-hoc neurobiotin and ankyrinG immunohistochemistry, which allowed us to define the location of the axon. APs were triggered by injecting incremental current steps, and the relationship between AP amplitude and AP HW was defined for each neuron before and after photo-ablation (**Figure 6B**). AP amplitude and AP HW were negatively correlated, and the slope of this relationship was modified by dendrite photo-ablation. Although the range of AP amplitudes was usually modified after photo-ablation, we could calculate the change in AP HW for APs of the same amplitude before and after dendrotomy (**Figure 6B, C**). Due to the severity of the experimental manipulation, this experiment could only be carried out on 7 neurons, including 3 nABD dendrotomies, 2 ABD dendrotomies, and 2 control neurons where laser illumination failed to induce dendrite photo-ablation. Consistent with the correlations described in **Figure 1** and the model predictions, nABD dendrotomy was associated with a decrease in AP HW (100 *vs* 89.4%, n=3), ABD dendrotomy was associated with an increase in AP HW (100 *vs* 113.65, n=2), while the absence of photo-ablation was associated with no change in AP HW (100 *vs* 99.1%, n=2; **Figure 6C**). These experiments strongly suggest that the opposite correlations between the topology of the ABD or nABDs and AP HW described in **Figure 1** arise because the ABD and nABDs have opposite causal influences on AP HW. Based on the modeling results (**Figure 5**), they also reinforce the hypothesis that sodium channel density is higher in the ABD than in nABDs.

**Figure 6.**
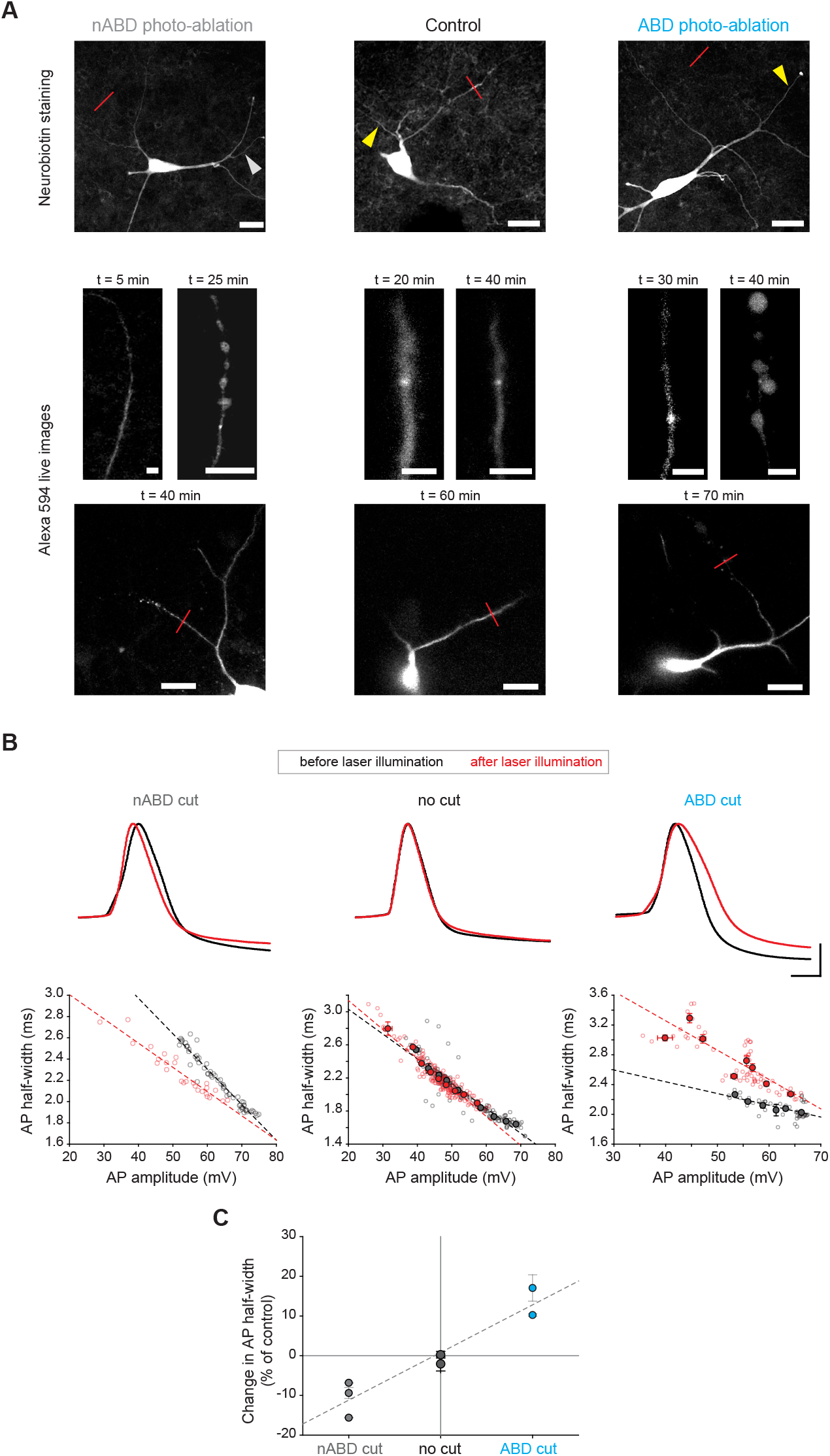
Dendrite photo-ablation confirms the opposite influence of nABDs and ABDs on AP half-width. **A**, **top row**, confocal images of post-hoc neurobiotin immunostaining of 3 neurons on which a dendritic photo-ablation experiment was performed (left, nABD dendrotomy; middle, no dendrotomy after laser illumination; right, ABD dendrotomy). The red line indicates where laser illumination (line scan) was performed, while the yellow arrowhead indicates the axon (identified by post-hoc ankyrinG immunostaining). **Middle row**, zoomed-in confocal live images showing the dendritic branch before laser illumination and after effective laser illumination for the 3 neurons presented in the top row images. **Bottom row**, zoomed-out confocal live images showing neuronal morphology after laser illumination. The exact location of laser illumination is indicated by a red line. **B**, AP waveforms recorded in the 3 neurons presented in panel **A** before (black traces) and after laser illumination (red traces). APs displaying a similar amplitude were chosen to allow comparison of the half-widths. **C**, scatter plots representing the relationship between AP amplitude and AP half-width before (black dots) and after laser illumination (red dots) in the same neurons presented in panels **A** and **B**. Light colors correspond to individual APs while binned and averaged values (with error bars, SEM) appear in dark color when the number of APs allowed averaging. Dotted lines indicate the linear regression depicting the relationship between AP amplitude and half-width. **C**, scatter plot summarizing the effect of dendrotomy observed in 8 neurons (3 nABD cut, 3 controls and 2 ABD cut). Scale bars: **B**, vertical 20mV, horizontal 2ms.

Somato-dendritic sodium channels underlie the faithful back-propagation of APs in the dendrites of SNc DA neurons (Hausser et al., 1995; Gentet and Williams, 2007; Seutin and Engel, 2010; Moubarak et al., 2019). While AP back-propagation is 100% reliable during pacemaking activity (Moubarak et al., 2019) or in response to long depolarizing steps of current (Gentet and Williams, 2007), numerous failures of back-propagation have been observed when APs were triggered by synaptic activity (barrages of EPSCs injected into the ABD or nABD) (Gentet and Williams, 2007). Specifically, APs failed to propagate efficiently to nABDs in particular when DA was applied onto the somato-dendritic compartment (Gentet and Williams, 2007). Since our results underlined the importance of dendritic topology and sodium channel density on the shape of the somatic AP, we wondered whether AP back-propagation could also be influenced by these same parameters. To test this, we simulated EPSC injection into the ABD in our multicompartment model, and recorded the triggered APs in the AIS and in the nABDs (**Figure 7A**). The same train of EPSCs was applied to all neurons in the center of the primary ABD, using either the homogeneous version of the model (model #1) or the heterogeneous version containing higher sodium and calcium conductances in the ABD (model #4). As indicated by their amplitude at the AIS recording site, all APs were initiated in the AIS and were considered to fail to back-propagate efficiently into the nABDs when the nABD voltage did not reach −15mV (**Figure 7B**). Consistent with the results presented in **Figures 3** and **5**, the percentage of back-propagation failures was negatively correlated with ABD complexity only in the heterogeneous model (r=-0.431, p=0.012, n=33, Pearson correlation coefficient) while the correlation was not significant in the homogeneous model (r=-0.146, p=0.417, n=33; the bootstrap analysis demonstrating that the difference in correlation between the two conditions is significant, p<0.003; **Figure 7C**). No relationship was found between the complexity or length of the nABDs and the percentage of failures (data not shown). These last results suggest that dendritic topology, and in particular ABD complexity, not only controls the shape of the AP at the soma but also its ability to faithfully back-propagate across the entire dendritic tree.

**Figure 7.**
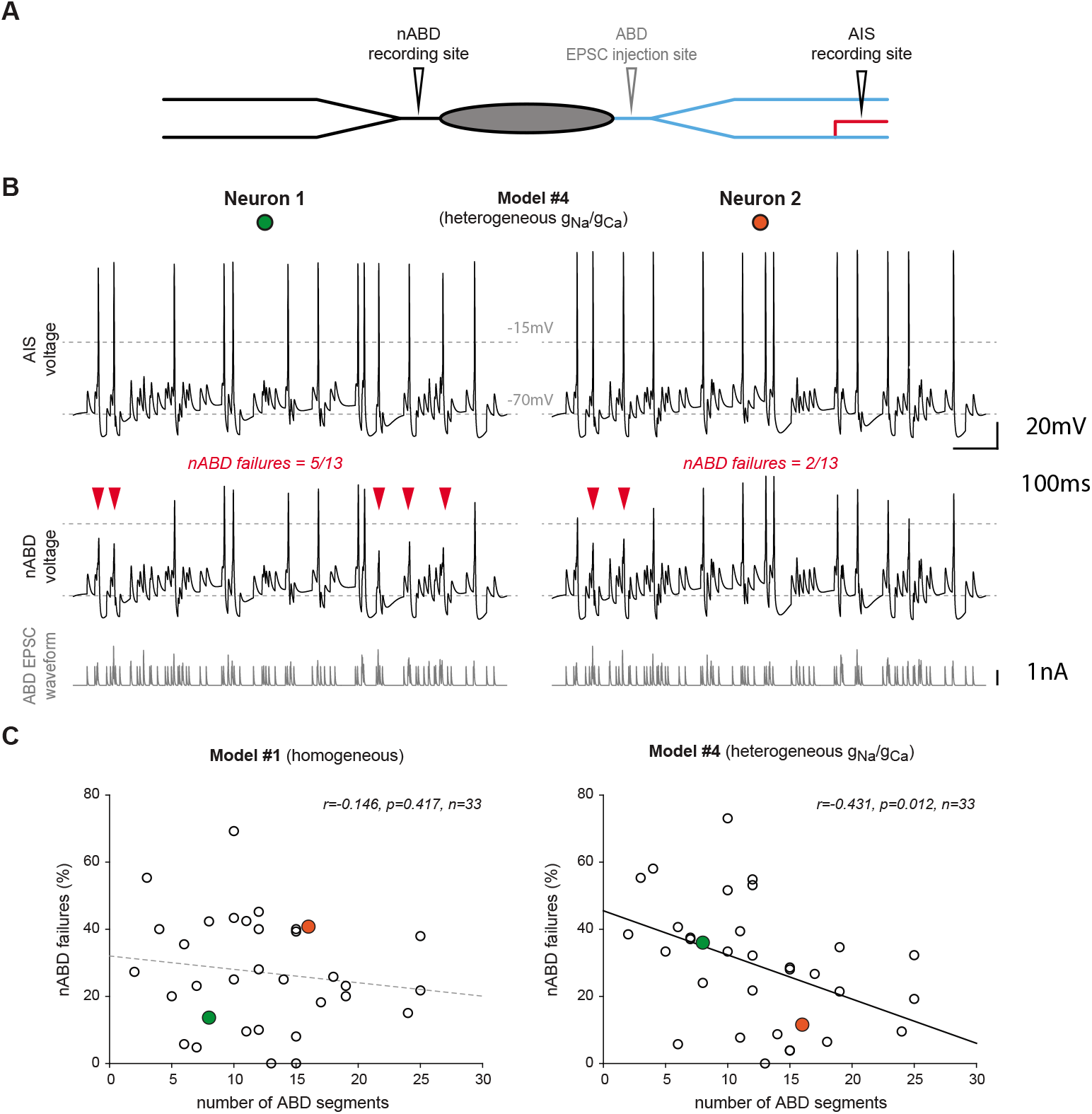
ABD complexity determines failures of AP back-propagation in response to barrages of EPSCs. **A,** schematic representation of the modeling paradigm. Barrages of EPSCs were injected into the ABD, while AP waveform was recorded in the AIS and nABD. **B,** voltage traces recorded in two different model neurons (using heterogeneous g_Na_/g_Ca_, model #4) in the AIS (top traces) and in the nABD (middle traces) in response to the same EPSC sequence injected in the ABD (bottom traces). AP back-propagation failures (indicated by red arrowheads) were detected as APs not reaching −15 mV (gray horizontal dotted line) in the nABD. **C,** scatter plots showing the relationship between ABD complexity and the percentage of nABD failures in the model #1 (homogeneous g_Na_/g_Ca_, left) and #4 (heterogeneous g_Na_/g_Ca_, right). The values corresponding to the two model neurons illustrated in panel **B** are identified by colored dots. A significant correlation (black plain regression line) was found only for the heterogeneous model (right scatter plot). Scale bars: **B,** vertical 20mV (upper traces), 1nA (lower traces), horizontal 100ms.

## DISCUSSION

In this study, we demonstrated that somatic AP shape in SNc DA neurons is strongly constrained by the specific topology of the dendritic arborization, due to the opposite influence of the ABD and nABDs on AP duration. Using computational modeling, we showed that this influence of dendritic topology on AP shape may require a higher density of sodium channels in the ABD. Moreover, in these conditions, AP back-propagation in response to synaptic activity also seems to be controlled by dendritic morphology, in particular ABD complexity.

The most important finding of our study is that the variation in dendritic topology in SNc DA neurons explains most of the cell-to-cell variation in half-width of the somatic APs (68% based on the multiple linear regressions presented in **Figure 1**). To our knowledge, this is the first experimental study demonstrating that dendritic morphology can have a predominant influence on AP shape across neurons of the same type, such that AP duration can be reasonably well predicted on the sole basis of dendritic topology. This finding is particularly surprising in the light of previous observations. Several studies have demonstrated that the biophysical properties of dendritic ion channels, including sodium channels, display substantial cell-to-cell variability in SNc DA neurons (Liss et al., 2001; Gentet and Williams, 2007; Seutin and Engel, 2010; Amendola et al., 2012; Engel and Seutin, 2015; Philippart et al., 2016; Moubarak et al., 2019). One may thus wonder whether their neuronal output is predominantly defined by variations in ion channel properties or in dendritic morphology. A previous study dedicated to the role of dendritic sodium channels in pacemaking showed that cell-to-cell variations in dendritic morphology have a greater influence on the spontaneous firing rate than variations in sodium channel density (Moubarak et al., 2019). The current study goes one step further by demonstrating that variations in dendritic morphology predict AP duration in spite of the substantial cell-to-cell variations in sodium channel density observed in this cell type. By showing that the relationship between dendritic topology and AP duration is reproduced when all neurons are endowed with identical sodium conductance density profiles, the computational modeling results reinforce these conclusions. Even so, we need to bear in mind that, at constant sodium conductance density, increasing dendritic complexity (and length) in the model leads to an increase in the overall sodium dendritic conductance. However, our goal was not to perform a sensitivity analysis comparing the relative efficiency of biophysical and morphological properties (Weaver and Wearne, 2008) in modifying AP shape, but rather to determine the minimal conditions required to explain the correlation observed between dendritic topology and AP duration. In this context, the most surprising result was that these conditions also enable the model to predict the duration of the AP recorded in real neurons, unambiguously demonstrating the predominant role of cell-to-cell variations in dendritic morphology in defining AP shape in this neuronal type.

Another interest of the current study is to shed light on the importance of morphology in defining AP shape. So far, most studies investigating the sources of variations in AP properties (duration, amplitude, energy efficiency) were focused on the contribution of the biophysical properties of sodium and potassium currents to these processes (Alle et al., 2009; Carter and Bean, 2009; Sengupta et al., 2010; Alle et al., 2011; Carter and Bean, 2011). Moreover, these studies aimed at understanding the biophysical basis of differences in AP shape across neuronal types and not within the same neuronal population. Briefly, large variations in AP duration across neuronal types seem to be essentially determined by the types of potassium channels expressed (Rudy and McBain, 2001; Bean, 2007; Carter and Bean, 2009). In particular, the expression of Kv3 and/or BKCa channels appears necessary to achieve fast AP repolarization and fast-spiking patterns of activity (Rudy and McBain, 2001; Lien and Jonas, 2003; Alle et al., 2011; Kaczmarek and Zhang, 2017; Hunsberger and Mynlieff, 2020). On the other hand, the variations in properties of sodium channels do not seem to contribute significantly to the differences in AP duration observed across cell types (Carter and Bean, 2009). Consistent with this, studies investigating the maturation of AP shape demonstrated that changes in duration during post-natal development in a given neuronal type are mainly related to changes in potassium channel expression patterns (Hunsberger and Mynlieff, 2020; Sanchez-Aguilera et al., 2020), although changes in sodium current have also been observed in some neuronal types (Dufour et al., 2014). One modeling study investigated specifically the influence of dendritic morphology on AP properties, focusing in particular on AP back-propagation in various neuronal types (Vetter et al., 2001). By comparing the back-propagating profiles in 8 different cell types, the authors demonstrated that dendritic morphology imposes strong constraints on the density of dendritic sodium channels necessary to achieve faithful back-propagation: neurons with a relatively simple dendritic tree such as SNc DA neurons require a fairly low density of sodium channels while Purkinje neurons never attain active dendritic AP propagation, even in the presence of a high sodium channel density (Vetter et al., 2001). More interestingly, these two cell types are less sensitive to changes in sodium channel density than most of the other neuronal types analyzed in the study (hippocampal and cortical interneurons and principal cells), which can display passive or active AP propagation depending on sodium channel density (Vetter et al., 2001). Our experimental and computational results extend these results by demonstrating that dendritic morphology also plays a critical role in defining the variation in AP shape across SNc DA neurons, in spite of substantial cell-to-cell variability in sodium channel density (Moubarak et al., 2019).

One may argue that the variability in ABD and nABD morphologies reported here is partly due to the slicing procedure, and that we may have overestimated the cell-to-cell variations in dendritic topology, due to artefactual sectioning of distal dendrites. While this may be true, the lack of a preferred orientation for the dendritic arborization of SNc DA neurons (Preston et al., 1981; Tepper et al., 1987; Lin et al., 2003) suggests that ABD and nABDs should have been affected with an equiprobability. Based on our results, such artefactual distal dendrotomy would mainly increase the range of variation of both dendritic complexities and AP duration, thus favoring the detection of a correlation between these parameters. Be that as it may, there is no reason to think that artefactual dendritic sectioning could create the correlation observed between AP shape and dendritic topology, and the results of our computational modeling and dendrotomy experiments provide biophysical explanations for this relationship.

An intriguing finding of the current study is that the statistical dependence between dendritic topology and AP HW is only reproduced when heterogeneous densities of g_Na_ and g_Ca_ are implemented in the computational model. In contrast, the outside-out recordings obtained in a previous study (Moubarak et al., 2019) did not reveal statistical differences in sodium current density between the ABD and the nABDs. This is not totally surprising since the method used to determine the patch area, and thus the capacitance, is based on an approximately linear but highly scattered relationship between pipette resistance ant patch area (Sakmann and Neher, 1995) (Figure 8, page 648; R^2^=0.15, p=0.01; our calculation). Although this method allows to evaluate the order of magnitude of current density, its prediction error may be as large as a factor four. Ion currents in DA neurons display a significant variability from neuron to neuron (Liss et al., 2001; Gentet and Williams, 2007; Seutin and Engel, 2010; Amendola et al., 2012; Engel and Seutin, 2015; Moubarak et al., 2019; Haddjeri-Hopkins et al., 2021) that would obscure the presence of subcellular heterogeneity when comparing g_Na_ between the ABD and nABDs in different neurons. Although outside-out recordings are widely used to estimate current densities, this technique is not sufficiently accurate to reveal a heterogeneous density of g_Na_ in the dendrites of SNc DA neurons in the range suggested by the model. In order to be consistent with our previous experimental results, we nonetheless tested the possibility that g_Na_ and g_Ca_ could be homogeneous within a same neuron but scale with ABD length across neurons (data not shown). Interestingly, while this manipulation reproduced the correlations between AP HW and dendritic topology, it failed to reproduce the opposite influences of ABD and nABD dendrotomies. On the other hand, increasing the excitability of the ABD compared to the nABDs by increasing g_H_ or decreasing g_K_ in the ABD failed to reproduce the correlation between AP HW and dendritic topology (**Figure 4B, C**), which necessitated specifically an increase in g_Na_ and/or g_Ca_. In conclusion, a slightly heterogeneous density of g_Na_ and g_Ca_ seems to be the simplest explanation of why ABD and nABDs have an opposite influence on AP HW.

A final important question is whether these opposite influences of ABD and nABDs, in addition to defining AP duration, have an impact on the physiology of SNc DA neurons. While the high density of somato-dendritic sodium channels participates in the robustness of pacemaking in these neurons (Tucker et al., 2012; Jang et al., 2014; Moubarak et al., 2019), it also supports the faithful back-propagation of APs (Hausser et al., 1995; Gentet and Williams, 2007; Seutin and Engel, 2010; Moubarak et al., 2019), which appears necessary for dendro-dendritic release of DA (Vandecasteele et al., 2008; Yee et al., 2018). Because of the axonal initiation of APs, gating the back-propagating APs would thus represent a way to decouple the axonal (striatal) and dendritic releases of DA: in fact Gentet and Williams (2007) demonstrated that somatic back-propagation failures occur in response to barrages of dendritic EPSCs and are potentiated by DA. Our modeling results confirm these observations and suggest that dendritic topology, in particular ABD complexity, is a determinant factor to predict the ability of an SNc DA neuron to efficiently back-propagate APs. Consistent with the other results presented in this study, this relationship only exists when the ABD contains a higher sodium conductance density than the nABDs. Thus, the specific dendritic topology of an SNc DA neuron would not only determine the duration of the AP at the soma, but also determine the robustness of AP back-propagation in response to barrages of excitatory synaptic inputs, therefore defining whether axonal and dendritic releases of DA are coupled or decoupled.

## ACKNOWLEDGMENTS

this work was supported by the French Ministry of Research (doctoral fellowship to E.M.) and the European Research Council (Consolidator grant 616827 *CanaloHmics* to J.M.G.). We thank Dr. D. Debanne for his support with the confocal live imaging experiments.

